# A plant lipocalin is required for retinal-mediated *de novo* root organogenesis

**DOI:** 10.1101/2020.11.09.375444

**Authors:** Alexandra J. Dickinson, Jingyuan Zhang, Michael Luciano, Guy Wachsman, Martin Schnermann, José R. Dinneny, Philip N. Benfey

**Affiliations:** Duke University, Durham NC; Carnegie Institute of Science, Stanford CA; Stanford University, Palo Alto CA; National Institutes of Health, Bethesda MD; Howard Hughes Medical Institute, Duke University, Durham, NC

## Abstract

Branching of root systems enables the exploration and colonization of the soil environment. In Arabidopsis roots, *de novo* organogenesis of lateral roots is patterned by an oscillatory mechanism called the root clock, which is dependent on metabolites derived from the β-carotene pathway^1, 2^. Retinoids are β-carotene-derived regulators of organogenesis in the animal kingdom. To determine if retinoids function in plant development, we conducted time-lapse imaging of a chemical reporter for retinoid binding proteins. We found that it oscillates with a comparable frequency to the root clock and accurately predicts sites of lateral root organogenesis. Exogenous application of retinal to wild-type plants is sufficient to induce root clock oscillations and lateral root organogenesis. A homology search yielded a potential *Arabidopsis* homolog, TEMPERATURE INDUCED LIPOCALIN (TIL) to vertebrate retinoid binding proteins. Genetic analysis indicates that TIL is necessary for normal lateral root development and a *til* mutant has decreased retinal sensitivity. TIL expression in a heterologous system conferred retinal binding activity, suggesting that it may directly interact with this molecule. Together, these results demonstrate an essential role for retinal and for plant retinal binding proteins in lateral root organogenesis.

---

Plants continuously develop post-embryonic lateral roots in order to forage for water and nutrients. The root clock, a temporal series of oscillating changes in gene expression, pre-patterns sites for initiation of lateral root primordia^3, 4^. Blocking carotenoid metabolism genetically or with inhibitors is sufficient to dampen root clock oscillations and prevent lateral root initiation^1, 2^. In vertebrate development, β-carotene-derived retinoic acid has important functions in the oscillatory somitogenesis clock^5^, neurogenesis, and vasculature development^6–8^. Because several carotenoid-derived metabolites have been shown to regulate root growth^2, 9–12^, we hypothesized that retinoids and their binding partners may function in the root clock.

Merocyanine aldehyde (MCA) is a chemical reporter designed for measuring vertebrate retinoid binding protein activity^13^. MCA has a Cyanine-5 (Cy5) head group fused to a retinaldehyde-like moiety that fluoresces when bound to a lysine residue in retinoid binding proteins (Figure 1A). To test for the presence of proteins that interact with retinal-like molecules in the root, we treated Arabidopsis plants with 10 μM MCA. The majority of the mature root remained non-fluorescent with MCA treatment. However, MCA-induced fluorescence exhibited striking spatial and temporal patterns in developing regions of the root (Movie S1, Figure 1B). Root meristems, regions of actively dividing cells, consistently displayed high levels of MCA fluorescence (Figure S1A). In the early differentiation zone, pulses of MCA fluorescence were observed (Figure 1B, Figure S1B), which were reminiscent of the root clock. To determine if MCA fluorescence pulses overlapped spatially or temporally with the root clock, we simultaneously tracked both processes using MCA treatment in roots expressing *pDR5:LUC*, an auxin-responsive reporter that also tracks the root clock. Spatially, MCA pulses overlapped with the “oscillation zone,” the region exhibiting maximal luminescence of the DR5 oscillatory expression (Figure S1C). Temporally, we found that the MCA pulse appeared, on average, 1.4 (+/− 0.9) hours after the DR5 expression maxima. These results suggest that MCA pulses may be triggered by the peak of the root clock as evidenced by a peak of DR5 expression. In addition, MCA-induced fluorescence in Arabidopsis roots suggests that one or more proteins capable of binding a retinal binding protein reporter have oscillatory properties and could be involved in the root clock.

**Figure 1.**
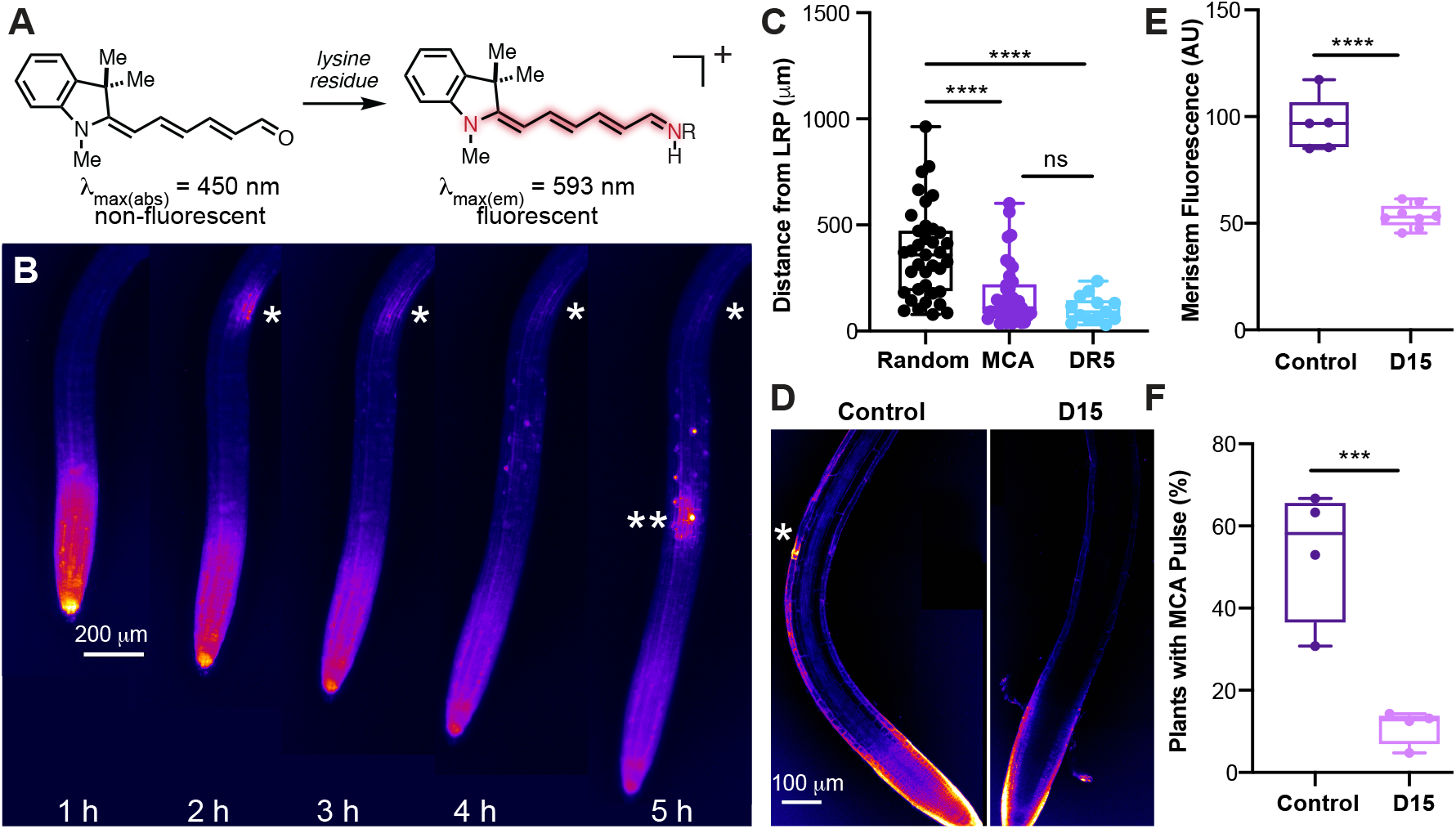
A reporter for retinal binding proteins predicts sites of lateral root organogenesis in Arabidopsis. A) The chemical structure and fluorescence properties of merocyanine aldehyde (MCA) in the absence and presence of a lysine residue from a retinal binding protein. B) Time course of MCA fluorescence dynamics in Arabidopsis roots. * and ** indicate the site of fluorescent oscillations in the early differentiation zone of the root. C) MCA oscillations predict sites of new lateral root primordia (LRP) as accurately as the DR5 root clock. The “random” control sites were selected using a random number generator that was linked to regions in the developing root. D) Confocal images of MCA fluorescence in control and D15 treated roots. * indicates the site of a fluorescent pulse in the root. E) Quantification of meristem fluorescence in control and D15 treated roots. F) Percent of roots with early differentiation zone MCA oscillations in control and D15 treated plants.

If MCA-binding protein activity is triggered by the root clock, then MCA pulses should correlate with lateral root primordia initiation. To test this hypothesis, we tracked the regions of the roots that experienced the maximum intensity of an MCA pulse over time. We identified early lateral root primordia 24-48 hours after the MCA pulse and found that, like the DR5 oscillations, MCA pulses accurately predict the sites of lateral root primordia (Figure 1C, Figure S2). This suggests that the activity of one or more putative plant retinal binding proteins is induced by the root clock and correlates with sites of lateral root specification.

To further test the connection between putative retinal binding protein dynamics and lateral root primordia development, we characterized MCA dynamics in roots in which DR5 oscillations are inhibited. D15 is a chemical inhibitor of carotenoid metabolism that reduces DR5 clock oscillations, blocks lateral root initiation, and decreases cell elongation^1, 2^. D15 had two major effects on MCA spatial and temporal patterning. First, it decreased MCA fluorescence in the meristem (Figure 1D, E). Second, it reduced the frequency of MCA pulses in the differentiation zone (Figure 1F). These results suggest that carotenoid metabolites are necessary for retinal binding in the root meristem and at sites of lateral root specification. Interestingly, D15 did not affect the amplitude of rare MCA pulses (Figure S3), suggesting that the effect of D15 on MCA pulses is binary – pulses either occur at full amplitude or they do not occur at all. These findings are consistent with a model in which the root clock oscillation triggers all or none retinal binding. Previous work demonstrates that apocarotenoid synthesis is necessary for initiating DR5 oscillations^1, 2^, suggesting that apocarotenoids may function at two important points in the root clock – before and after the DR5 maxima. Importantly, these data are consistent with the hypothesis that retinal binding proteins are involved in lateral root formation.

The correlation between MCA dynamics and lateral root development suggests that retinal, or a structurally similar compound, could be important for lateral root organogenesis (Figure 2A, Figure S4A). To determine if retinoids naturally occur in Arabidopsis and are affected by D15 treatment, we analyzed extracts from plants treated with D15 or a mock control using HPLC-MS. We found four compounds present in Arabidopsis that are decreased in D15-treated plants: retinal (apo16), 14’-apo-beta-carotenal (apo14), 12’-apo-beta-carotenal (apo12), and 10’-apo-beta-carotenal (apo10) (Figure S4A-B). To determine if retinal, apo14, apo12, or apo10 could rescue D15 inhibition of the lateral root clock, we treated D15-inhibited roots with these compounds and quantified resulting changes in DR5 oscillations. We found that retinal, apo12, and apo14 significantly increased the amplitude of DR5 oscillations in D15-inhibited roots (Figure S4C). 1 μM of retinal added to D15-treated roots was sufficient to fully rescue the amplitude of root clock oscillations within 24 hours (Figure 2B-C, Movie S2). At this concentration, retinal also rescued cell elongation in D15 inhibited roots (Figure S4D). We next monitored the ability of retinal, apo12, and apo14 to rescue D15 inhibition of lateral root development. Only retinal and apo14 fully rescued D15 inhibition of primary root growth and lateral root capacity, which is defined as the ability of plants to form lateral roots after the primary root has been excised (Figure 2D-E). Retinal and apo14 significantly increased the number of lateral root primordia in uninhibited plants (Figure 2F), indicating that exogenous application of either retinal or apo14 is sufficient to induce ectopic organogenesis. Exogenous retinal treatment increased lateral root density (Figure S4E) as well as the total number of root primordia. These findings reveal that retinal and apo14 are sufficient to induce lateral root organogenesis. Notably, it is possible that apo14 is metabolized to retinal, and that retinal or another retinoid is the active signaling compound required for lateral root development.

**Figure 2.**
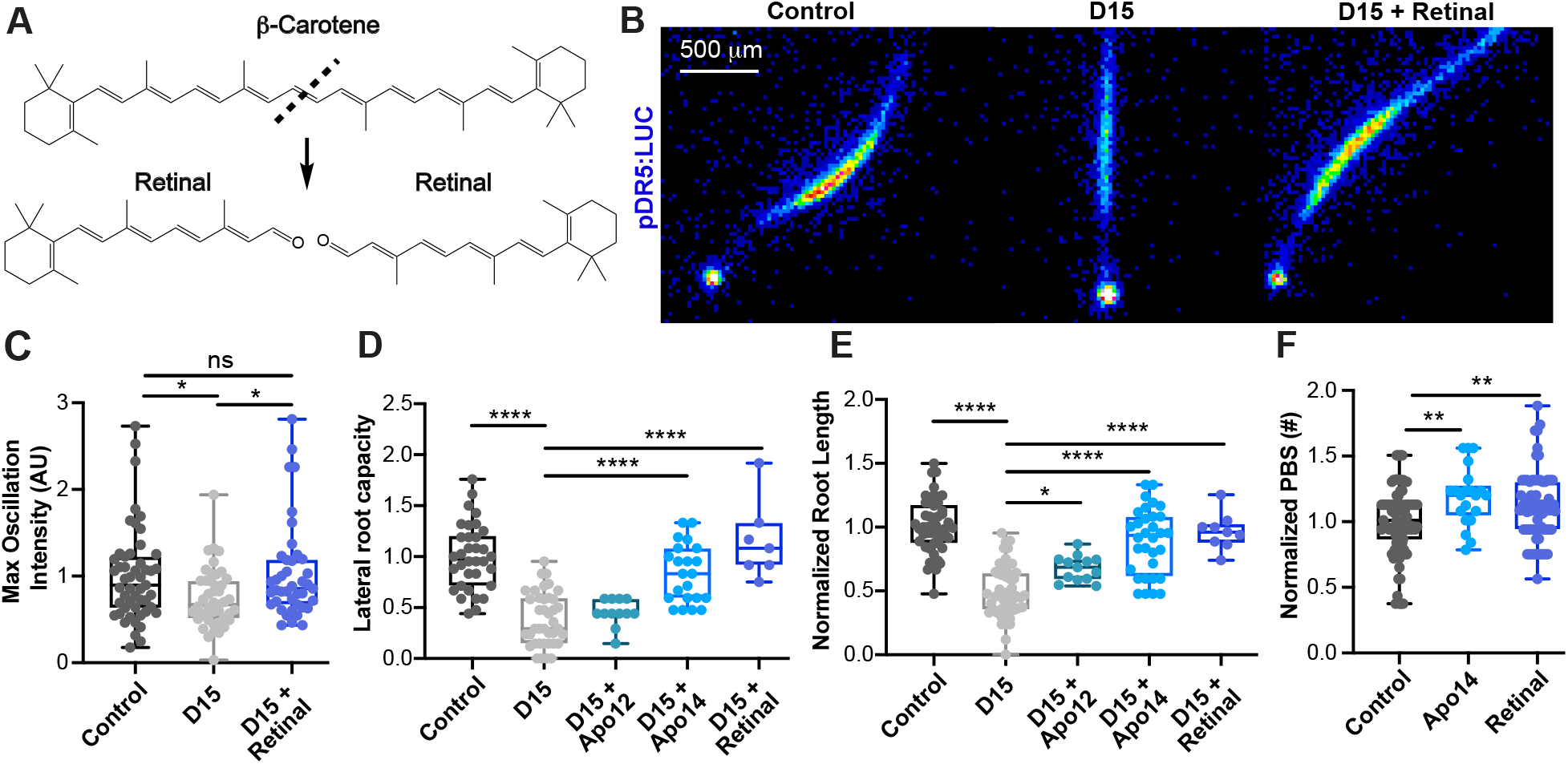
Retinal rescues D15 inhibition of the root clock and lateral root organogenesis. A) Chemical structures of retinal and its precursor, β-carotene. B) Luminescent images of the maximum root clock oscillation in pDR5:LUC roots with the following treatments: vehicle control, D15, and D15 in combination with retinal. C) Quantification of the maximum root clock oscillation intensity in pDR5:LUC roots. D) Quantification of the capacity of plants to form lateral roots in D15 inhibited roots (normalized to the vehicle control). E) Root length in plants exposed to D15 (normalized to the vehicle control). F) The number of lateral root pre-branch sites in pDR5:LUC roots treated with apo14 or retinal (normalized to the vehicle control).

We searched for Arabidopsis proteins with sequence homology to retinal binding proteins, such as opsins, from algae and vertebrates, but did not find any. However, we did identify homology between vertebrate and plant lipocalins, cross-kingdom protein transporters of small, hydrophobic molecules^14, 15^. TIL (AT5G58070), the primary lipocalin expressed in the Arabidopsis root, has sequence homology (E-value = 9e-04, % identity = 25%) and predicted structural homology (TM-score = 0.72) to RETINOL BINDING PROTEIN 4 (RBP4), a vertebrate lipocalin (Figure 3A-B, Figure S5). TIL has been previously implicated in preventing lipid peroxidation, particularly during light and heat stress^16^. Analysis of an existing RNA-seq expression map of the root during DR5 oscillations revealed that TIL is expressed during the root clock^17^. In these experiments, roots were dissected into five regions during the DR5 oscillation maxima: the region rootward of the oscillation (“r”), the oscillation region (“p”), the region directly shootward of the oscillation (“s”), the most recently formed pre-branch site (“pbs”) marking the cells undergoing organogenesis, and the region immediately shootward of the new pre-branch site (“ss”). TIL was most highly expressed in the regions shootward of the oscillation zone (Figure 3C). Wachsman *et al.* also dissected the root DR5 oscillation zones at three time points: before the DR5 oscillation maxima (“bp”), during the oscillation maxima (“p”), and after the oscillation maxima (“ap”). TIL expression was highest after the peak of the DR5 clock (Figure 3D).^17^ This gene expression pattern is similar to that of the MCA fluorescence oscillations, which are strongest following the DR5 maxima, suggesting that it is a good candidate for a plant retinal binding protein.

**Figure 3.**
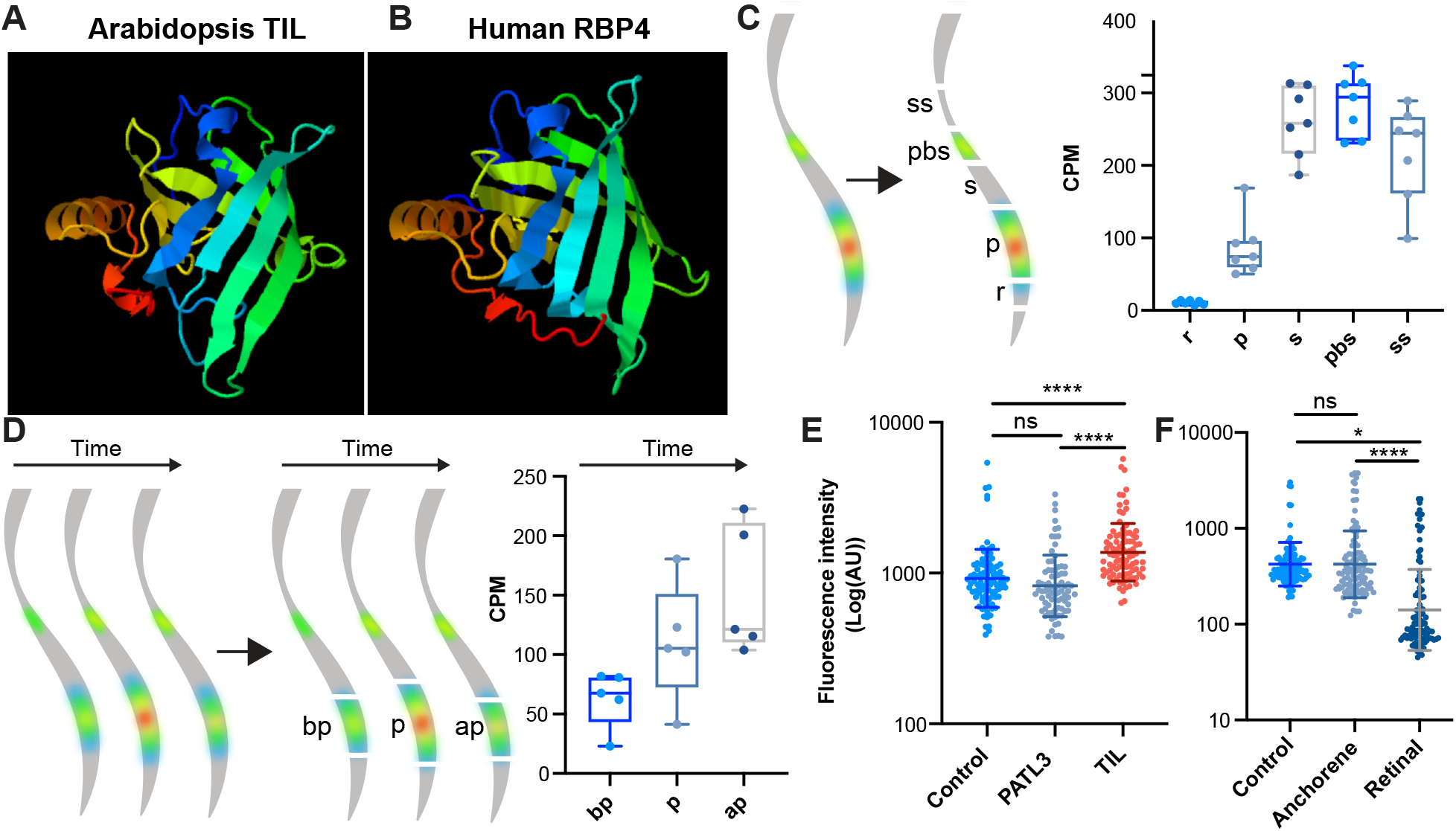
TIL is a plant lipocalin that interacts with retinal. A-B) Predicted protein structure of Arabidopsis TIL and Human RBP4 using the protein homology/analogy recognition engine V 2.0 (PHYRE2)^19^. C) Gene expression (in counts per million) of TIL in different regions of the root undergoing DR5 oscillations^17^. D) Gene expression (in counts per million) of TIL during root clock oscillations.^17^ Root schematics are adapted from Wachsman *et al*^17^. E) Fluorescence intensity of MCA treated *E. Coli* transformed with genes from Arabidopsis. The control cells were not transformed. F) Fluorescence intensity of MCA treated *E. Coli* transformed with TIL and pre-treated with a DMSO control, anchorene, or retinal.

To test if TIL binds retinal, we heterologously expressed TIL in *E. coli* and measured the effect on MCA fluorescence. As a control, we expressed PATL3, a plant lipid binding protein with a cellular retinaldehyde binding protein (CRAL)-TRIO domain. PATL3 expression did not change MCA fluorescence, indicating that the ability to bind lipids and homology to retinal binding protein domains are not sufficient to induce MCA fluorescence (Figure 3E, Figure S6). TIL significantly increased MCA fluorescence, indicating that it interacts with MCA (Figure 3E, Figure S6). Next, we hypothesized that if TIL binds to retinal, application of retinal should disrupt TIL-induced MCA fluorescence. Application of retinal to TIL-expressing E. coli significantly reduced MCA fluorescence, indicating that retinal and MCA compete for TIL binding (Figure 3F, Figure S7). In contrast, cells treated with anchorene, a carotenoid-derivative that is chemically similar to retinal, did not diminish MCA fluorescence. These results suggest that TIL interacts with retinal but not to a related apocarotenoid.

To determine the role of TIL in organogenesis, we examined root development in previously characterized TIL mutant alleles^16^. Because both *til-1* and *til-2* mutant alleles had similar significant reductions in lateral root branching (Figure 4A-B), we focused further experimentation on the *til-1* allele. The *til-1* mutant had a defect in root growth due to a reduction in cell elongation, which phenocopies inhibition of carotenoid biosynthesis with D15 (Figure 4C). To determine if TIL was important for establishing the root clock or for later stages of primordia development, we examined the expression of DR5 in *til-1*. We did not observe changes in frequency or intensity of the DR5 oscillation in *til-1*, suggesting that TIL is not necessary for root clock oscillations. Instead, we found that *til-1* mutants are slower to initiate lateral root primordia after the peak of the DR5 oscillation (Figure 4D, Figure S8, Movie S3). This supports a model in which retinal synthesis is necessary for both DR5 and MCA oscillations (Figure S9) and TIL plays a post-DR5 role that links the root clock to the divisions that form lateral root primordia. Overall, these results indicate that TIL is critical for proper lateral root organogenesis and that loss of TIL function leads to defects consistent with reduced retinal signaling.

**Figure 4.**
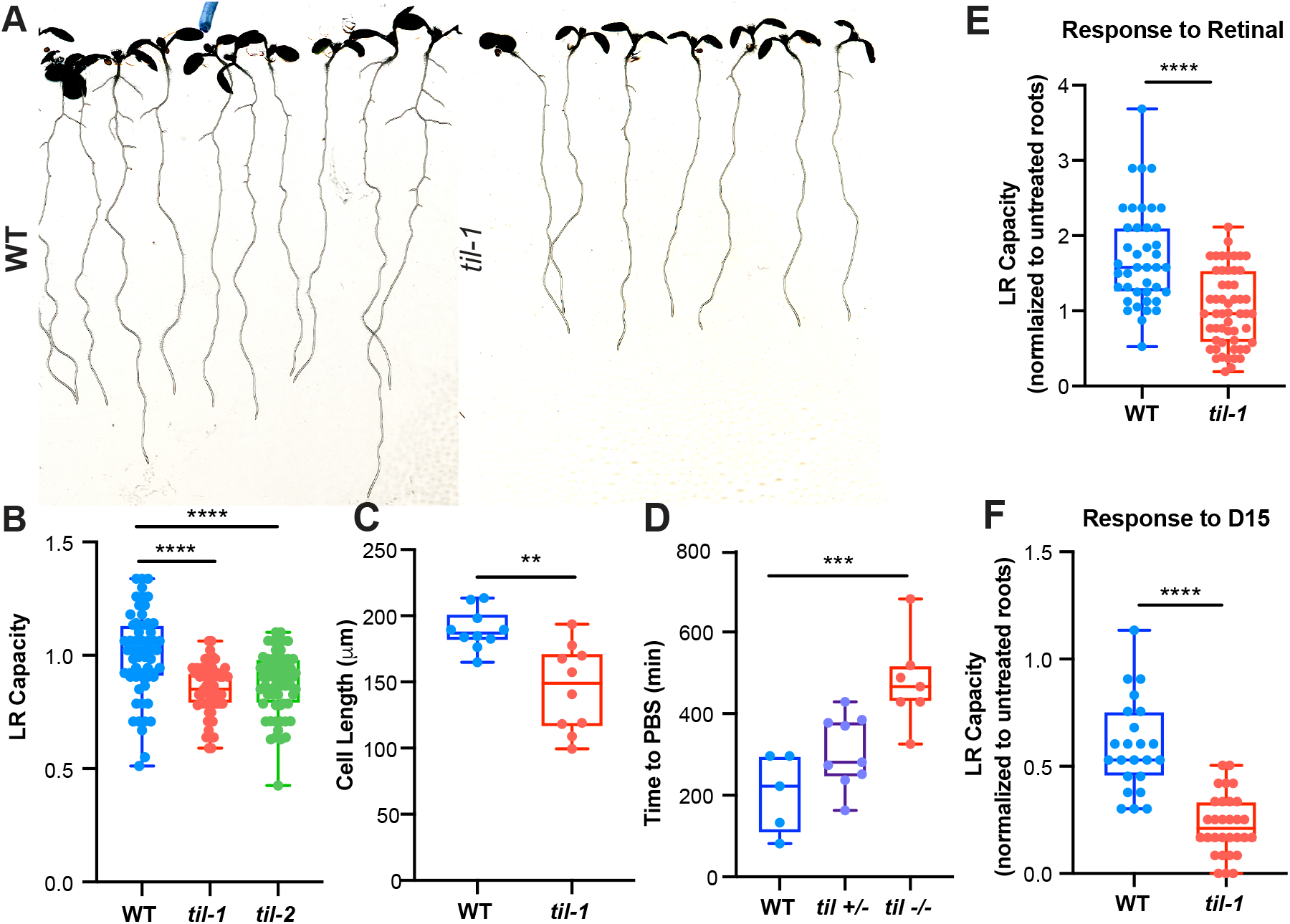
TIL is essential for proper lateral root development. A) Images of WT and *til-1* mutant seedlings. B) Lateral root capacity (LRC) in WT and *til* mutants (normalized to WT roots). C) Average length of mature cortex cells in WT and *til-1* plants. D) The time between the root clock maxima and the formation of a lateral root pre-branch site, measured in pDR5:LUC roots. E-F) Fold change in lateral root (LR) capacity in response to retinal (E) and D15 (F). Values are normalized with respect to the untreated roots for each genotype. Genotype by treatment interactions were significant (≤ 0.003).

To explore the relationship between the plant lipocalins and the retinal response, we treated *til-1* roots with both retinal and D15 and quantified the effects on lateral root development. We found that *til-1* roots were less sensitive to retinal, indicating that retinal-induced root organogenesis is dependent on TIL (Figure 4E). In contrast, D15 treatment caused an increase in inhibition of lateral root capacity in *til-1* mutants, suggesting that these mutants are more sensitive to D15 than WT plants (Figure 4F). This indicates that reducing retinal biosynthesis in *til-1* further aggravates the mutant phenotype, suggesting that additional retinal perception pathways are still functional in *til-1* mutants. Overall, the changes in sensitivity in *til-1* provide genetic evidence that TIL is involved in retinal-mediated effects on lateral root development.

We identified retinal as an endogenous metabolite that is sufficient to induce post-embryonic root organogenesis. Reduction of retinal biosynthesis leads to inhibition of the root clock, the first known stage of lateral root organogenesis. Oscillations of activity of a chemical retinal binding protein reporter predicted sites of lateral root organogenesis. We identified TIL, a plant lipocalin able to bind retinal, and found that it is involved in converting the root clock oscillation into an initiated root primordium. To date, at least three pathways have been identified that play a role in regulating the root clock. In addition to apocarotenals, these are pectin modification^17^ and auxin^18^. It will be interesting to determine if these pathways act independently, or in interlocking processes. In addition, these results open up new questions about the convergent use of retinal related compounds in the regulation of developmental clocks across the plant and animal kingdoms.

## Materials and Methods

### Plant Growth and Treatment Conditions

Detailed experimental procedures are provided in the Supplemental Materials. All *Arabidopsis thaliana* plants were in the Columbia-0 background. Null mutants *til-1* (SALK_136775C) and *til-2* (SALK_150259C) were obtained from the ABRC. Retinal, apo-14, and apo-12 were generously supplied by Earl Harrison (Ohio State University). Apo-10 was generously supplied by Loredana Quadro (Rutgers). Working concentrations of retinal and apo-14 for long term (> 24 hour) experiments were 1 μM and 500 nM, respectively. For long term experiments, plates were wrapped in parafilm and the roots were grown on treated media and kept in the dark. For these experiments, the shoots were not directly treated with media and were the only part of the plant exposed to light. The merocyanine aldehyde (MCA) used is related to a previously described compound^13^, see supplemental information for a full description of the synthesis and characterization of the aldehyde and putative chromophore formed.

### Root Phenotyping

Merocyanine aldehyde (MCA) was synthesized as previously described (Yapici). To measure MCA fluorescence in plants, roots were treated with 10 μM MCA and imaged using dsRED filter sets. Fluorescence was tracked over time via stereomicroscopy or confocal microscopy. Lateral root capacity, or the ability of the plant to produce lateral roots, was measured as described previously^1^. Briefly, the root meristems were excised 5 days post-germination. This induces lateral root emergence. Three days after excision, the total number of emerged lateral roots was quantified. Lateral root clock oscillations were characterized using time-lapse imaging as previously described by Moreno-Risueno, *et al*. Roots were sprayed with 5 mM potassium luciferine (Gold Biotechnology) and then were imaged every 7 min over the course of 18 h using a chemiluminescence imaging system (Roper Bioscience).

### Chemical Analysis

HPLC-MS samples were prepared by homogenizing whole seedlings 5 days post germination in liquid nitrogen. Apocarotenoids were extracted from 100-200 mg of homogenized root tissue using propanol with 0.1% butylated hydroxytoluene. LC-MS/MS was performed using an Agilent 1260 HPLC coupled to an Agilent 6520 Q-TOF ESI mass spectrometer. Separation was conducted in an a 5 μm, 2 × 100 mm Gemini NX-C18 column (Phenomenex). The mobile phase solvents used were water with 0.1% (v/v) formic acid (Solvent A) and acetonitrile with 0.1% (v/v) formic acid (Solvent B). A linear gradient with a flow rate of 0.4 mL/min was used over the course of the separation. The mobile phase gradient used was 50:50 (Solvent A:Sovlent B) to 3:97 (Solvent A:Solvent B) over 12 minutes. The qTOF MS parameters were set as follows: mass range, 50-1000 m/z; gas temperature, 350 C, drying gas flow rate, 11 L/min, nebulizer, 35 psig.

### *E. coli* Assays

MCA fluorescence in *E. coli* was measured as described previously (Yapici). Briefly, *E. Coli* was transformed with a plasmid allowing LacI-inducible expression of an empty vector (control), PATL3, or TIL. Cells at OD_600_ were treated with IPTG and grown for 6 hours at 37° C. Cells were treated with 10 μM merocyanine for 30 minutes, and then imaged using a dsRED imaging set up on a fluorescence microscope. For apocarotenoid binding experiments, cells were incubated with IPTG for 6 hours at 37° C, then pre-treated with 200 μM retinal or 200 μM anchorene for 30 minutes. Following this, cells were treated with 10 μM merocyanine and then imaged.

### Statistical Analysis

For all experiments with more than two samples, data was analyzed using one-way ANOVAs with Tukey’s multiple comparison tests. All experimental analyses have a one-way ANOVA p value ≤ 0.05. For experiments with two samples, data was analyzed using unpaired t-tests. The symbols *, **, ***, and **** indicate p values ≤ 0.05, 0.01, 0.001, and 0.0001, respectively.

**Supplementary Information** is available for this paper.

## Acknowledgements

We thank Dominique Bergmann, Miguel Moreno-Risueno, Loredana Quadro, and Isaiah Taylor for their critical reading of the paper and insights. We thank Elizabeth Sattely for use of her lab equipment and Medhavinee Mijar for her technical support during this project. This work was supported by the Howard Hughes Medical Institute and the US National Institutes of Health (MIRA 1R35GM131725 (to PNB) and by the Arnold and Mabel Beckman Postdoctoral Fellowship (to AJD). The research of JRD was supported in part by a Faculty Scholar grant from the Howard Hughes Medical Institute and the Simons Foundation. ML and MS are supported by the Intramural Research Program of the National Institutes of Health (NIH), the National Cancer Institute, and the Center for Cancer Research.

## Author contributions

AJD and PNB conceived of the work and drafted the manuscript. AJD, PNB, JZ, ML, GW, MS, and JRD designed the experiments and interpreted the data. AJD, JZ, ML, and GW acquired the data. All authors revised the manuscript.

