## Supporting Information for "A plant lipocalin is required for retinal-mediated *de novo* root organogenesis"

**Supplemental Information**

**A plant retinal-binding protein is essential for *de novo* root organogenesis**

### Synthesis and Characterization of Merocyanine Aldehyde (MCA)

**General Materials and Methods.** Unless stated otherwise, reactions were conducted in oven-dried glassware under an atmosphere of nitrogen or argon using anhydrous solvents (passed through activated alumina columns). All commercially obtained reagents were used as received. Flash column chromatography was performed using silica on a CombiFlash® Rf 200i (Teledyne Isco, Inc.). High-resolution LC/MS analyses were conducted on a Thermo-Fisher LTQ-Orbitrap-XL hybrid mass spectrometer system with an Ion MAX API electrospray ion source in negative ion mode. Analytical LC/MS was performed using a Shimadzu LCMS-2020 Single Quadrupole utilizing a Kinetex 2.6 μm C18 100 Å (2.1 x 50 mm) column obtained from Phenomenex, Inc. Runs employed a gradient of 0→90% MeCN/0.1% aqueous formic acid over 4.5 min at a flow rate of 0.2 mL/min. $^{1}$H NMR and $^{13}$C NMR spectra were recorded on Bruker spectrometers (at 400 or 500 MHz or at 100 or 125 MHz) and are reported relative to deuterated solvent signals. Data for $^{1}$H NMR spectra are reported as follows: chemical shift (δ ppm), multiplicity, coupling constant (Hz), and integration. Data for $^{13}$C NMR spectra are reported in terms of chemical shift. Absorption curves were performed on a Shimadzu UV-2550 spectrophotometer operated by UVProbe 2.32 software. Fluorescence traces were recorded on a PTI QuantaMaster steady-state spectrofluorimeter operated by FelixGX 4.2.2 software, with 5 nm excitation and emission slit widths, 1.0 s integration rate, and emission correction enabled. Data analysis and curve fitting were performed using MS Excel 2016 and GraphPad Prism 8.

**Synthetic Procedures**

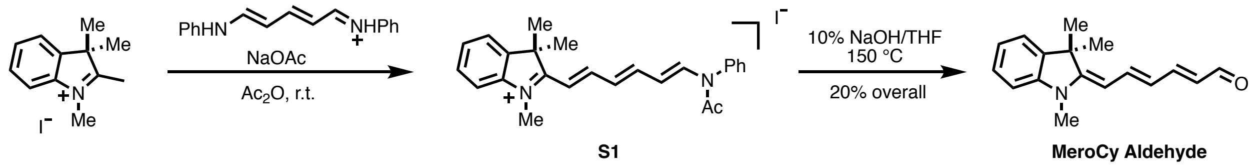


**Intermediate S1.** 1,2,3,3-Tetramethyl-3H-indolium iodide (301 mg, 1 mmol), glutaconaldehydedianil hydrochloride (284 mg, 1 mmol, 1 equiv.), NaOAc (172 mg, 2.1 mmol, 2.1 equiv) and acetic anhydride (12 mL) were added to a scintillation vial equipped with a magnetic stir bar. The reaction mixture was stirred at room temperature for 0.5 h. When the starting material was consumed (reaction control: LC-MS), the reaction mixture was divided into two portions and precipitated into $Et_2O$ (30 x 2 mL) in two falcon tubes. The dark purple suspension was centrifuged for 5 min at 4500 RPM and the supernatant decanted. Each pellet was resuspended in $Et_2O$ (30 mL), the centrifugation process was repeated, and the pellets were dried under vacuum for 1 h (< 0.1 Torr). The crude product was purified by automated flash chromatography (40 g silica, 0 $\rightarrow$ 15% MeOH/$CH_2Cl_2$) to afford the known intermediate **S1** as a purple solid in 66% yield (331 mg) with spectroscopic data in agreement with that reported in the literature.[1] $^1$H NMR (400 MHz, $CDCl_3$) δ 8.17 (d, $J$ = 13.7 Hz, 1H), 7.85 (dd, $J$ = 15.0, 11.3 Hz, 1H), 7.62 – 7.45 (m, 8H), 7.22 – 7.15 (m, 3H), 6.90 (dd, $J$ = 14.4, 11.2 Hz, 1H), 5.41 (dd, $J$ = 13.7, 11.4 Hz, 1H), 4.22 (s, 3H), 1.97 (s, 3H), 1.72 (s, 6H) ppm; $^{13}$C NMR (100 MHz, $CDCl_3$) δ 180.66, 169.51, 156.06, 151.82, 142.65, 141.66, 141.24, 138.02, 130.84, 130.21, 130.04, 129.56, 128.99, 128.23, 122.51, 114.04, 113.79, 113.64, 77.48, 77.16, 76.84, 51.57, 36.09, 27.24, 23.48 ppm.

**MeroCy Aldehyde.** Intermediate **S1** (331 mg) was dissolved in THF (6.7 mL) and treated with 10% NaOH (20 mL) and heated to 150 °C in a sealed microwave vial for 18 h, during which time the reaction mixture changed color from purple to yellow. The reaction mixture was allowed to cool and was extracted with

---

[1] Berezin, M. Y.; Guo, K.; Teng, B.; Edwards, W. B.; Anderon, C. J.; Vasalatiy, O.; Gandjbakhche, A.; Griffiths, G. L.; Achilefu, S. *J. Am. Chem. Soc.* **2009**, *131*, 9198-9200.

$CH_2Cl_2$ (3 x 20 mL). The combined organic layer was dried over $Na_2SO_4$, concentrated by rotary evaporation, and purified by automated flash chromatography (12 g silica, 0 → 50% EtOAc/hexanes) to afford **MeroCy Aldehyde** in 20% yield (50 mg) as an orange-brown solid. $^1$H NMR (400 MHz, DMSO-d6) δ 9.39 (d, *J* = 8.3 Hz, 1H), 7.54 – 7.43 (m, 2H), 7.31 (d, *J* = 7.3, 1.2 Hz, 1H), 7.19 (t, *J* = 7.7, 1.2 Hz, 1H), 6.94 – 6.84 (m, 2H), 6.23 (dd, *J* = 13.9, 11.5 Hz, 1H), 5.91 (dd, *J* = 14.7, 8.3 Hz, 1H), 5.55 (d, *J* = 12.5 Hz, 1H), 3.19 (s, 3H), 1.55 (s, 6H) ppm; $^{13}$C NMR (100 MHz, DMSO-d6) δ 192.38, 162.33, 155.50, 144.17, 141.84, 138.86, 127.79, 124.61, 121.69, 121.42, 120.53, 107.29, 96.46, 45.86, 40.15, 39.94, 39.73, 39.52, 39.31, 39.10, 38.89, 29.08, 27.81 ppm; HR-MS (ESI) calculated for $C_{17}H_{19}NO$ $(M+H)^+$ 254.1539, observed 254.1530.

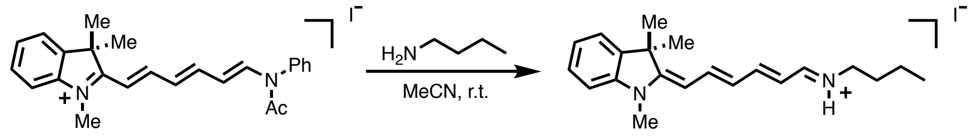

S1 MeroCy-*n*Bu

**MeroCy-*n*Bu. S1** (9.0 mg, 0.018 mmol) was dissolved in dry MeCN (200 μL) in a 1-dram vial equipped with a magnetic stir bar and the solution was flushed with nitrogen. *n*-butylamine (11.9 μL, 0.12 mmol, 5 equiv.) was added under nitrogen and the reaction mixture was stirred for 5 minutes, during which time the color changed from red to magenta and LC-MS analysis indicated full conversion to the *n*-butylamine adduct. The reaction mixture was evaporated by nitrogen stream, and the crude residue was purified by automated flash chromatography (4 g silica, 0 → 15% MeOH/$CH_2Cl_2$) to afford **MeroCy-*n*Bu** in 87% yield (6.3 mg) as a magenta film. The $^1$H NMR spectrum of **MeroCy-*n*Bu** indicates the presence of a mixture of isomers, as described for a similar compound.[2] See below for optical characterization.

**Determination of Molar Absorption Coefficients**

[2] Herwig, L.; Rice, A. J.; Bedbrook, C. N.; Zhang, R. K.; Lignell, A.; Cahn, J. K. B.; Renata, H.; Dodani, S. C.; Cho, I.; Cai, L.; Gradinaru, V.; Arnold, F. H. *Cell Chem. Biol.* **2017**, *24*, 415-425.

Molar absorption coefficients (ε) were determined in MeCN using Beer's law, from plots of absorbance vs. concentration. Measurements were performed in 10 mm path length quartz cuvettes (Hellma 111-QS), maintained at 25 °C, with absorbance at the highest concentration ≤ 1.0.

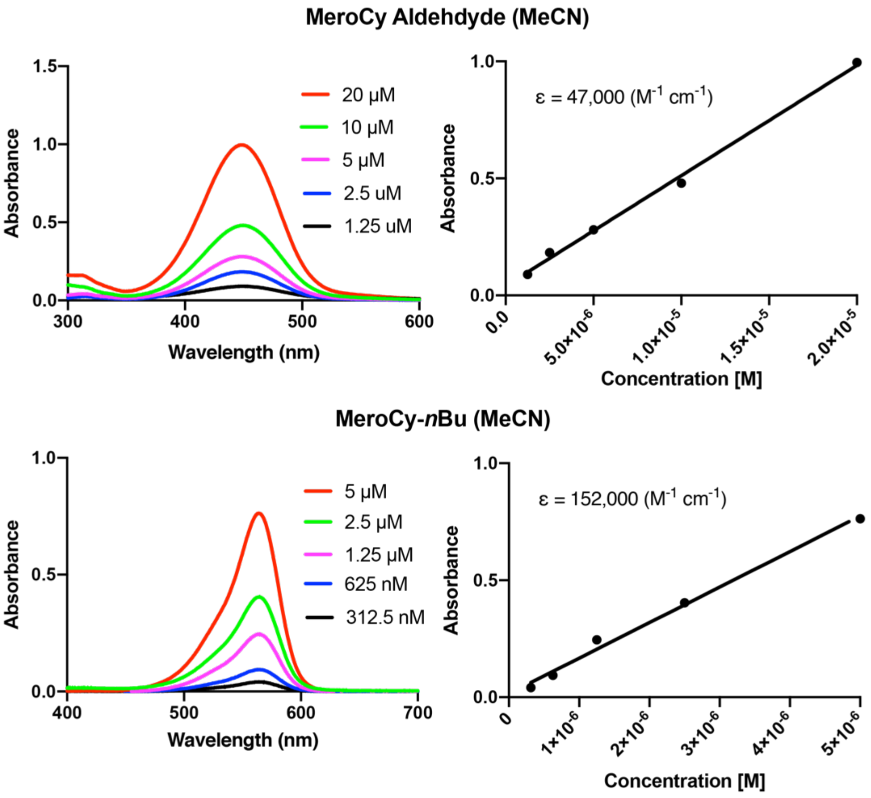


Left: Absorbance spectra of the compounds indicated in MeCN. Right: Absorbance values at $\lambda_{max}$ as a function of concentration. The slope of the line corresponds to the molar absorption coefficient ε.

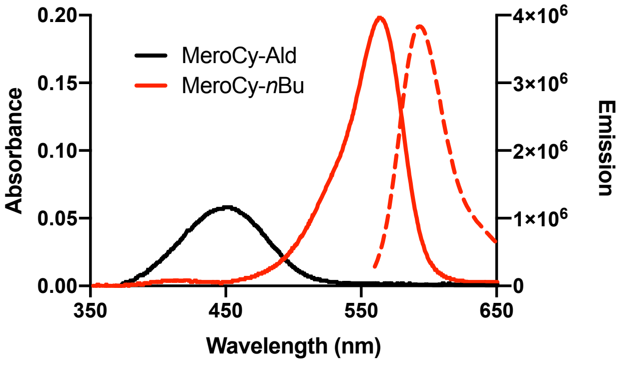


Absorbance (solid lines) and emission spectra ($\lambda_{excitation}$ = 520 nm, dashed line) of the compounds indicated at a concentration of 1 µM in MeCN.

### Supplementary Movies

Movie S1: MCA fluorescence dynamics in a growing root. The scale bar is 200 µm. The duration of the movie is 15 hours. The images in this movie have been corrected for photobleaching using the ImageJ Histogram Matching algorithm.

Movie S2: The root clock in *pDR5:LUC* plants treated with D15. One hour before imaging, roots were treated with a vehicle control (left) or retinal (right). The scale bar is 0.5 mm. The duration of the movie is 8 hours.

Movie S3. The *pDR5:LUC* root clock and pre-branch site formation in WT and *til-1* plants. The root on the left is WT and the root on the right is *til-1*. The scale bar is 0.5 mm. The duration of the movie is 13.5 hours.

### Supplementary Figures

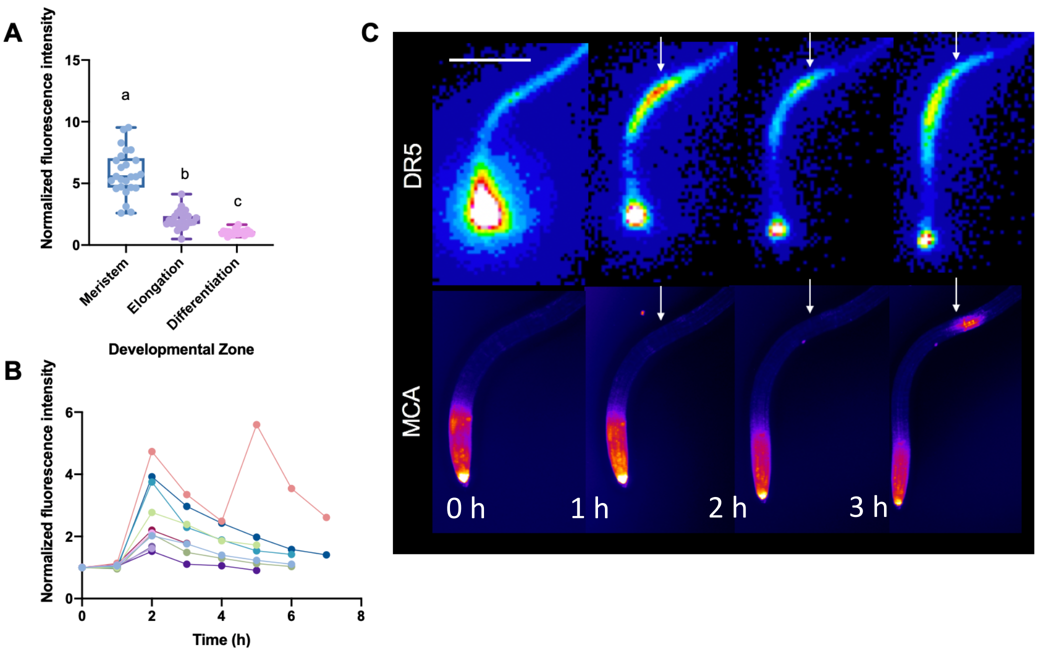


Figure S1. Merocyanine aldehyde (MCA) treatment of Arabidopsis roots reveals spatial and temporal dynamics of plant retinoid binding proteins. A) MCA fluorescence intensity in root meristems, elongation zones, and differentiation zones that are not experiencing an MCA oscillation. B) MCA fluorescence intensity during MCA pulses in the elongation/differentiation zones. The time frame of the oscillations was normalized to show all the oscillations synchronously (the peak of the oscillation was set to 2 hours for each root). C) Images of a *pDR5:LUC* root over time treated with MCA. The top row contains luminescence images. The bottom row contains fluorescence images of the same root.

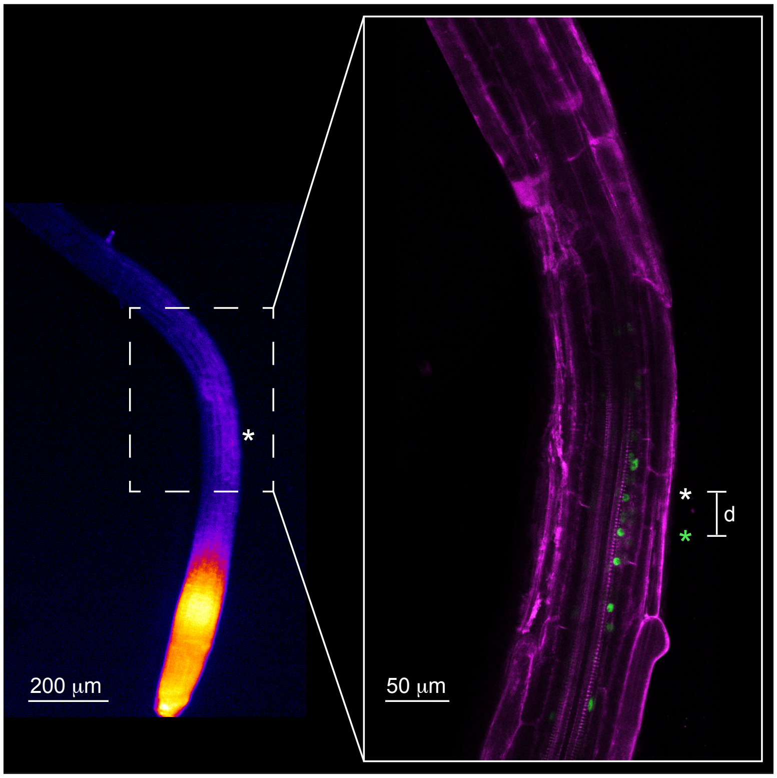


Figure S2. Tracking MCA pulses over time in *pGATA23:GFP* roots[3]. In order to characterize the physical relationship between the site of MCA oscillations and lateral root primordia, we tracked roots over the course of several days using fluorescence microscopy. The white asterisks indicate the sites of MCA oscillation maxima. The green asterisk indicates the center of a lateral root primordium, identified two days after MCA imaging. The distance between the site of maximal MCA fluorescence and the center of the new lateral root primordium was measured to determine how accurately the MCA pulse predicted the location of the lateral root primordium. All MCA and lateral root primordia sites were chosen without referencing each other. The "random" control site was chosen using a random number generator that was linked to a position in the root.

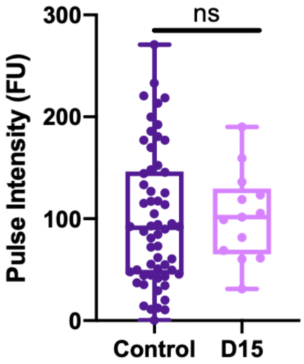


[3] De Rybel, B.; Vassileva, V.; Parizot, B.; Demeulenaere, M.; Grunewald, W.; Audenaert, J.; Van Campenhout, J.; Overvoorde, P.; Jansen, L.; Vanneste, S.; Möller, B.; Wilson, M.; Holman, T., Van Isterdael, G.; Brunoud, G.; Vuylsteke, M.; Vernoux, T.; De Veylder, L.; Inzé, D.; Weijers, D.; Bennett, M. J.; Beeckman, T. *Current Biology* **2010**, *19*, 1697-1706.

Figure S3. Fluorescence intensity of MCA pulses in control and D15 treated roots.

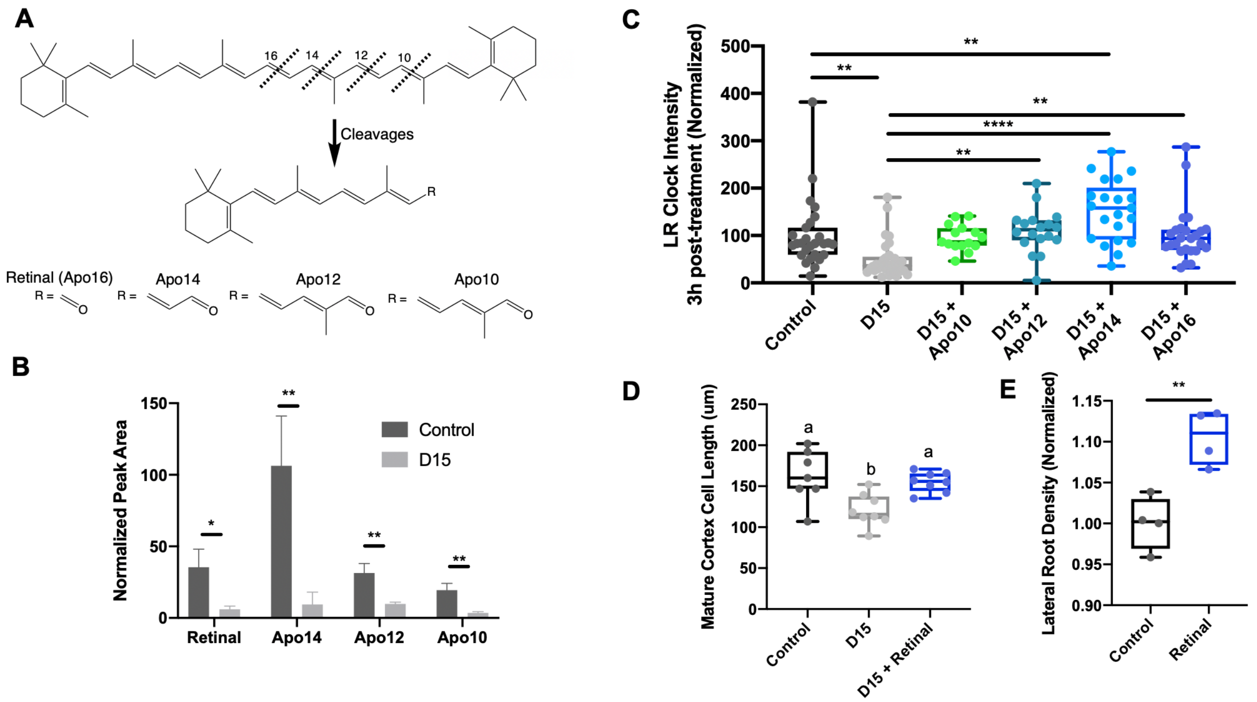


Figure S4. Certain apocarotenals can rescue D15 treatment phenotypes. A) Chemical structures of beta-carotene and its apocarotenal cleavage products produced by cleaving the double bonds before the 10th and 16th carbons. B) Relative amount (as measured by HPLC chromatogram peak area) of four apocarotenals in control and D15 treated plants. Peak area was normalized to the peak area of linoleic acid in each sample. C) Maximum intensity of the root clock as measured by DR5 expression in plants treated with D15 and apocarotenals. D) Cell length of mature cortex cells in plants treated with D15 and retinal. E) Lateral root density in control and retinal treated plants (the points shown are the averages of four experiments).

**A**

| Score | Expect | Method | Identities | Positives | Gaps |
|---|---|---|---|---|---|
| 24.3 bits(51) | 9e-04 | Compositional matrix adjust. | 35/142(25%) | 55/142(38%) | 30/142(21%) |

```
Query  7    MEVVKGLNVERYMGRWYEIASFP--SRFQPKNGVDTRATYTLNPDGTIHVLNETWSNGKR  64
             +V +   +  RY G WY +A         F    N V    A +T++ +G +         + G+
Sbjct  30   FKVKENFDKNRYSGTWYAMAKKDPEGLFLQDNVV---AQFTVDENGQMSAT----AKGRV  82

Query  65   GFIEGSAYKAD------PKSDEAKLKVKFY-VPPFLPIIPVTGDYWVLYIDPDYQHALI-  116
                      AD           D AK  K+K++  V   FL         D+WV   +D DY      +
Sbjct  83   RLFNNWDVCADMIGSFTDTEDPAKFKMKYWGVASFLQ--KGNDDHWV--VDTDYDTYALH  138

Query  117  ---------GQPSRSYLWILSR  129
                      G   + SY  ++ SR
Sbjct  139  YSCRELNEDGTCADSYSFVFSR  160
```

**B**

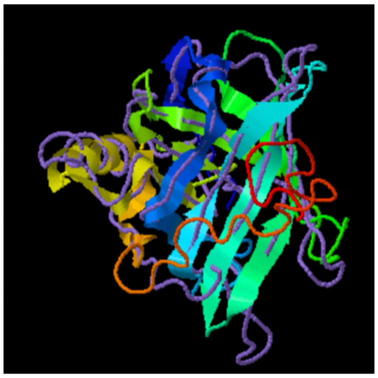

Figure S5. Protein homology between *A. thaliana* TIL and *G. gallus* RBP4. A) Sequence alignment procured using the National Center for Biotechnology Information (NCBI) protein Basic Local Alignment Search Tool (BLAST). B) Overlay of predicted protein structures of *A. thaliana* TIL (rainbow) and *G. gallus* RBP4 (purple) using iTASSER.

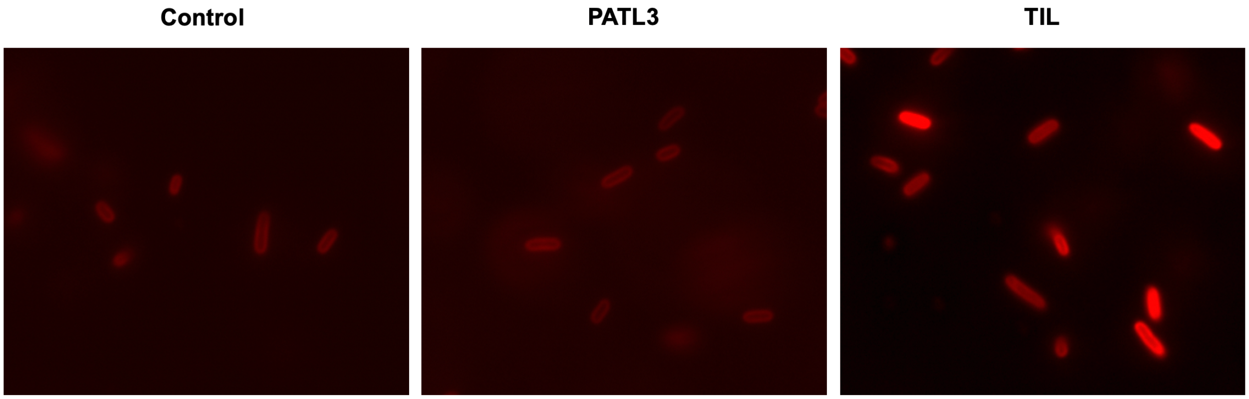


Figure S6. Fluorescence imaging of *E. coli* treated with MCA. In the left panel, control 5-alpha *E. coli* cells (New England BioLabs) are not expressing any *A. thaliana* genes. In the middle panel, cells are expressing *A. thaliana* PATL3. In the right panel, cells are expressing *A. thaliana* TIL.

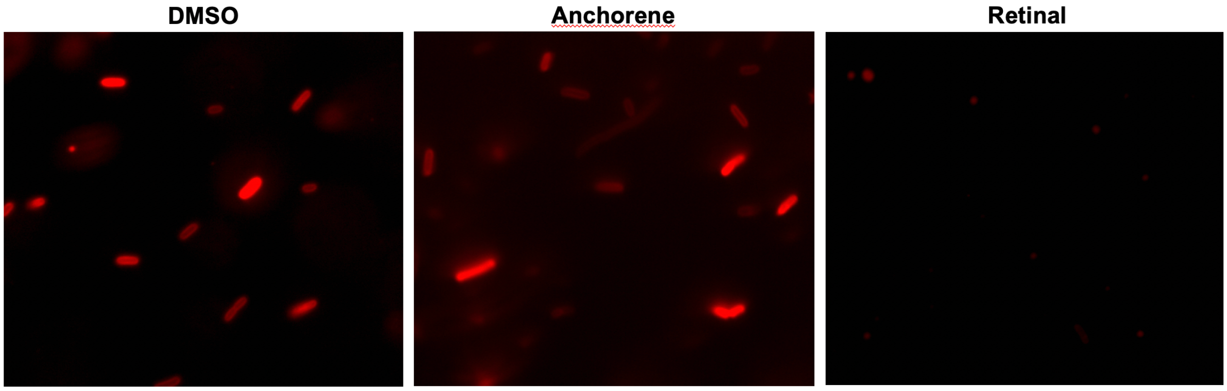


Figure S7. Fluorescence imaging of *E. coli* expressing *A. thaliana* TIL co-treated with MCA and DMSO, anchorene ((2*E*,4*E*,6*E*)-2,7-dimethylocta-2,4,6-trienedial), or retinal.

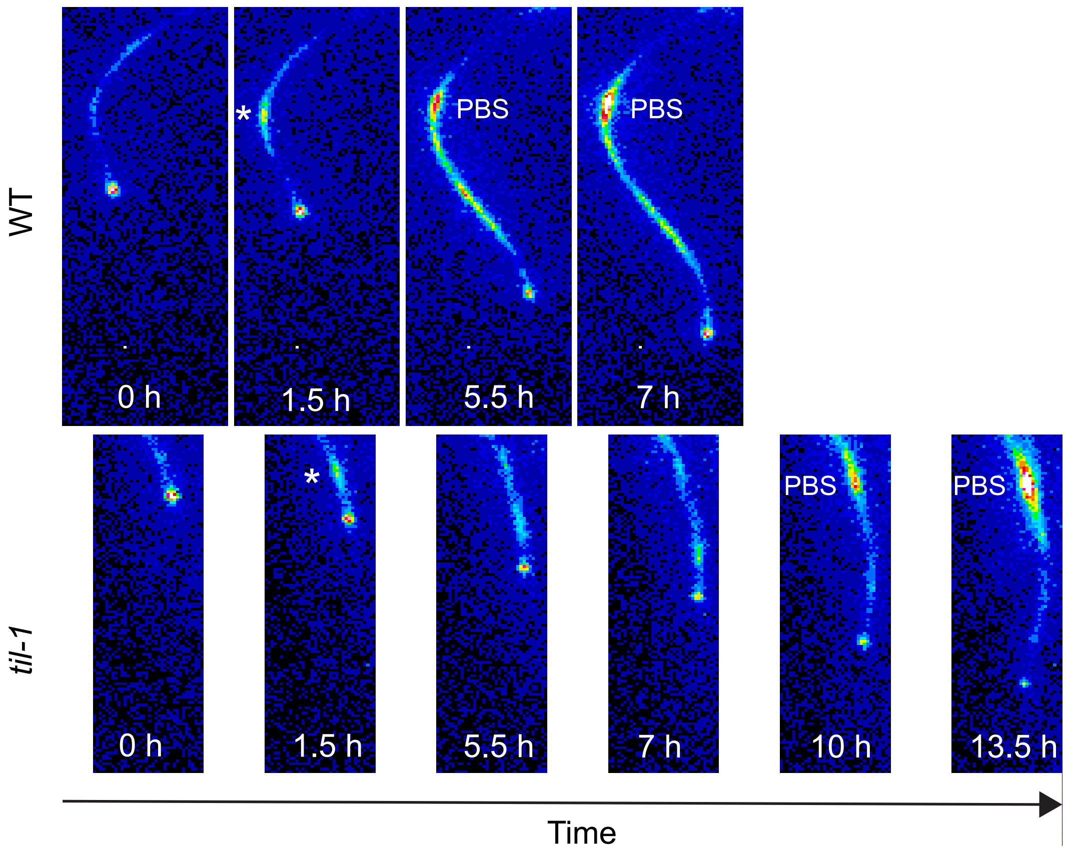


Figure S8. Time course of *pDR5:LUC* root clock oscillations in WT and *til-1* (see Movie S3 for full time course). * marks the maximum DR5 oscillation, which was normalized to occur at 1.5 hours for each root. The first timepoints when the corresponding pre-branch sites (PBS) were clearly visible are shown for each root. In WT, this occurred at 5.5 h. In *til-1*, this occurred at 10 h.

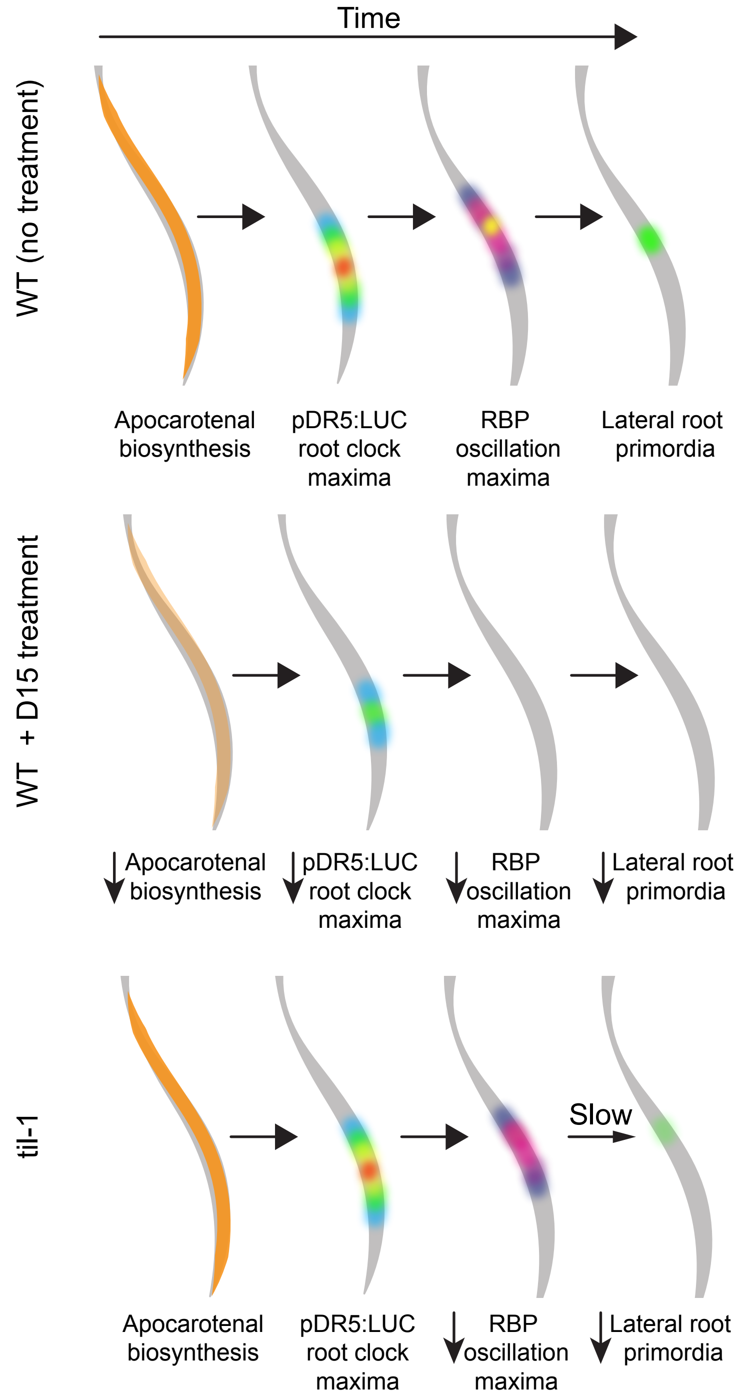


Figure S9. Schematic showing the relationship between synthesis of the retinal and apo14 apocarotenals, DR5 clock oscillations, plant retinoid binding protein (RBP) activity, and lateral root primordia pre-branch sites.
